# The Relationship Between Motor Fitness, Fundamental Movement Skills, and Functional Movement Screen in Primary School Children

**DOI:** 10.1101/2020.08.04.235879

**Authors:** Hua Wu, Wichai Eungpinichpong, Hui Ruan, Xinding Zhang, Xiujuan Dong

## Abstract

This study investigates motor fitness (MF), fundamental movement skills (FMS), and functional movement screen (FMS™) in 7-10-year-old children, and evaluates the relationship between FMS , MF and FMS™ combination with Seefeldt’s model for empirical research, thus to present effective strategies of physical development in children. A class was randomly selected in four school grades (1-4) along with 30 students from each to take the Test of Gross Motor Development-2 (TGMD-2). A total of 117 children (42 girls, aged 7-10) participated in three tests: TGMD-2, MF tests, and FMS™. MF levels (good, fair, and poor) and FMS™ levels (high, moderate, and low) were classified according to specific percentile ranges. A multiple (R×C) chi-square test analysis of the relationship between MF, FMS, and FMS™ was applied and post hoc testing estimated the possibility of FMS and FMS™ predicting MF. The results showed that only 43% of children were rated “good” on MF. Most fourth-grade students exhibited a certain gap with mature FMS (TGMD-2 score 70.13±9.68< 96 full scores). Boys scored significantly higher on the object control subtest and the TGMD-2 total score compared to girls (p<0.001), while girls had a significantly greater score than boys on the FMS™ (*p*=0.001). The results of multiple chi-square demonstrated FMS to be weakly correlated with MF, χ^2^ (4,N=117) =14.605, p =0.006< 0.01, Cramer’s V = 0.25. Both 60.5% of “excellent” FMS and 59.6% of “high” FMS™ children were categorized as having a “good” MF level. On the other hand, only 23.1% of the “worst” FMS and 24.3% of “low” FMS™ individuals were classified as having a “good” MF level. Our results suggest that MF, FMS, FMS™ are relatively independent systems linking with each other, generating mutual interaction in children’s motor development. At different stages or different advantages of them motor development, we may emphasize training one or a few parts.

## Introduction

The role of fundamental movement skills (FMS) has increasingly attracted attention from scholars, being recently considered part of the primary pedagogical approach for children.[1,2] FMS has been defined as a common activity with specific patterns, locomotion (such as running, hopping and sliding), and object-control (such as dribbling, throwing, kicking).[3] Previous studies show that FMS correlates with physical activity and health-related physical fitness in children and teenagers,[4–7] and low FMS proficiency has been found in children, [1,8] with the prevalence rarely above 50%,[9] and the vast majority performed below average.[10]

Seefeldt [11] first presented the role of FMS in life-long physical activity in a hypothetical model, which suggested that incompetency in FMS created a hypothetical “proficiency barrier” for an individual to attain motor proficiency (such as sports and games). In his model, reflexes serve as the basis for all future movements, so FMS are built upon a base of reflexes. The importance of the FMS phase in motor development models is also highlighted in the ‘mountain model’[12] and ‘clock hourglass model.’[13] Clearly, from these viewpoints and experiences with children aged 6-18 years,[14,15], teaching FMS seems to be a significant foundation for motor development and physical activity.

All three models suggest that reflexes and/or rudimentary movements provide a neurological basis for high level movement development. Functional Movement Screen(FMS™) tests [16] were developed based on fundamental proprioceptive and kinesthetic awareness principles, discovered through observation of an infant’s growth and motor development and suggesting proprioception which performs basic motor tasks by reflexes. However, successful movement patterns first formed during early childhood (such as squatting and the flexibility of limbs), may worsen due to a variety of factors, including sedentary behavior, training, and age.

Since then, the FMS^™^ has become a popular intra-rater and inter-rater reliability tool[17,18] to analyze the ability of athletes to perform certain basic movements.[19] It has also been used in children to access functional fitness,[20] to evaluate the relationship between FMS^™^, children’s weight status and physical inactivity,[21,22] as well as athletic performance.[23]

Motor development models are applied in sports practice, and these tests are widely used, but lack of empirical researches on associations among FMS, motor fitness (MF), and FMS™ , especially have yet to be properly investigated in children, which may influence research on strategies to promote physical activity and athletic development in children. Our study has two purposes: (a) to examine the MF, FMS, and FMS^™^ level in children aged 7-10 years, and (b) to explore the association between MF, FMS, and FMS^™^, especially to explore the relationship of FMS competence and FMS^™^, to levels of MF.

## Materials and Methods

### Subjects

This study used a cross-sectional design involving the relationship between MF, FMS, and FMS.^™^ We used a statistical program to randomly assign four classes from different grades (grade1 to grade 4) in a primary school (Haikou) in China, including 249 healthy children aged 7–10 years. The children first took the MF test[24] and the Test of Gross Motor Development-2 (TGMD-2). Then, from TGMD-2 scores, 120 children were selected as a sample (30 children from each class) to undergo the FMS^™^ test (Fig 1). Finally, 117 children underwent the three tests, including the MF test extracted from Chinese National Student Physical Fitness Standard, [24] TGMD-2[25], and FMS^™^ .[16] Table 1 demonstrates the descriptive statistics of each grade group. Participant and parental consent were obtained before testing and the study was approved by the university ethics committee. The study also conformed to the declaration of Helsinki.

**Fig 1.**
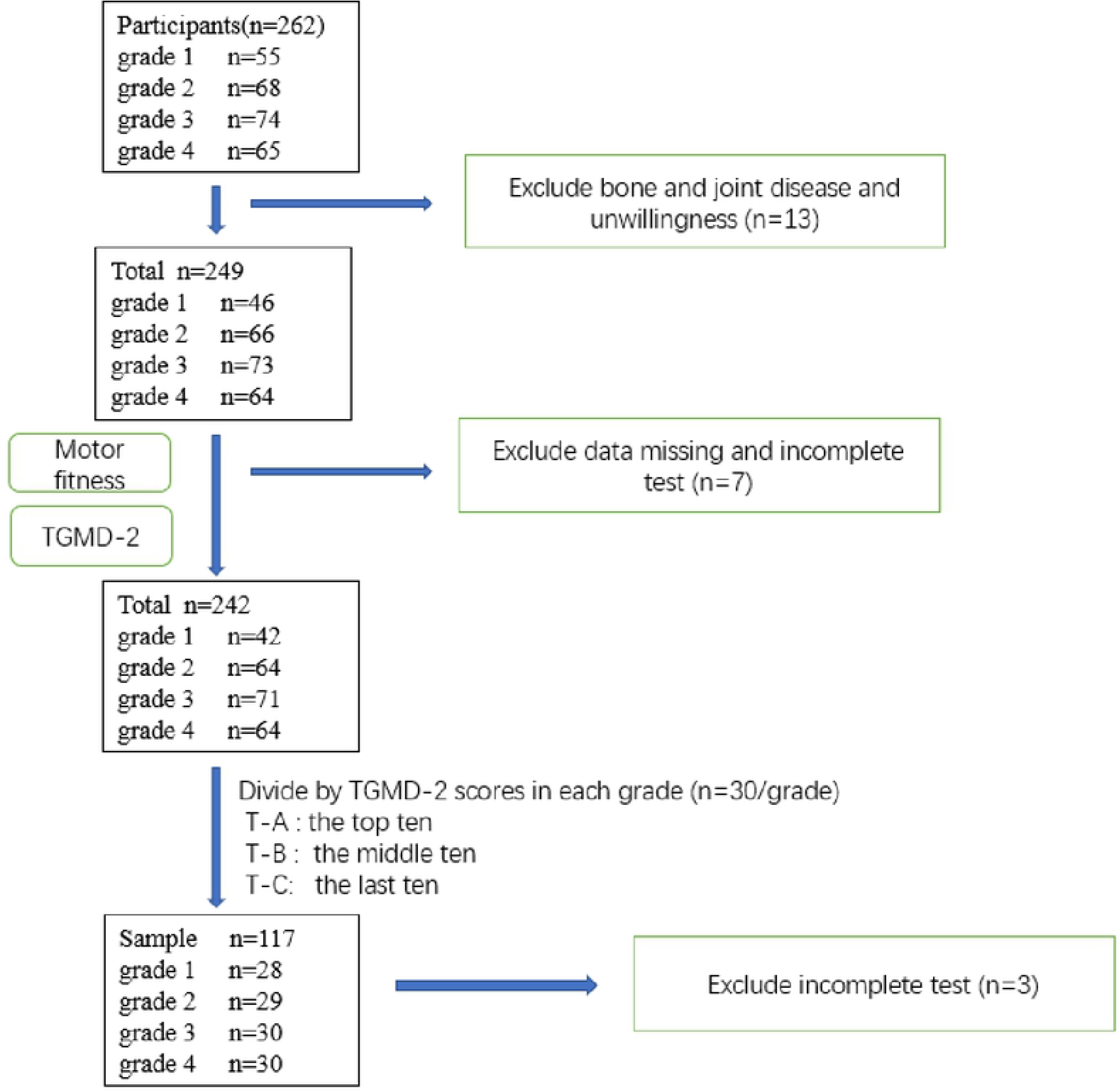
Flow chart showing the number of the sample selection process.

**Table 1.** Descriptive statistics for each grade group (mean ± SD).

| Group | Boys |  |  | Girls |  |  |  |
| --- | --- | --- | --- | --- | --- | --- | --- |
|  | Height (cm) | Weight (kg) | n | Height (cm) | Weight (kg) | n | total |
| **Grade 1** | 122.30±5.15 | 24.61± 5.32 | 19 | 119.97±4.90 | 21.68±3.34 | 9 | 28 |
| **Grade 2** | 130.22±5.88 | 28.12±7.46 | 17 | 129.54±4.53 | 26.12±4.48 | 12 | 29 |
| **Grade 3** | 135.19±4.85 | 31.03±4.53 | 20 | 130.86±3.81 | 26.80±5.01 | 10 | 30 |
| **Grade 4** | 137.23±5.88 | 31.59±6.14 | 22 | 133.75±8.01 | 31.00±10.61 | 8 | 30 |

### Measurement Procedure

Before the tests (except FMS^™^), participants finished warming-up which consisted of jogging followed by joint exercise of 2 minutes. After the children familiarized themselves with the test rules, they completed the tests following the Fig 1 protocol. The interval between each test was of three days. The test was conducted during normal school hours. Raters and data collectors were chosen by their physical education teachers and qualified testers.

### MF test

To compare the sport ability in motor development models, we selected part of the 2014 revised Chinese National Student Physical Fitness Standard (CNSPFS)[24,26] as our test. Grade 1 and 2 students completed a 50m sprint, sit and reach, and timed rope-skipping as part of the test, while timed sit-ups were added to the three exercises mentioned above for students from grade 3-4 (Table 2). Speed, flexibility, coordination, and strength are the components of MF which are needed for success in athletics and lifetime sport and activities.[27] Following the scoring criteria, raw data were converted into scores. The final total score was weighted by test scores, where 70 was considered a perfect score.

**Table 2.** MF indicators and weight coefficient on CNSPFS for testing the students of different grades.

| School grade | MF indicators | Objective | Weight score |
| --- | --- | --- | --- |
| **Grade 1 and**<br>**Grade 2** | 50 m sprint | Speed of movement | 20 |
|  | Sit and reach | lower-back/upper-thigh flexibility | 30 |
|  | Timed rope-skipping | coordination | 20 |
| **Grade 3 and**<br>**Grade 4** | 50 m sprint | Speed of movement | 20 |
|  | Sit and reach | lower-back/upper-thigh flexibility | 20 |
|  | Timed rope-skipping | coordination | 20 |
|  | Timed sit-ups | Abdominal strength and endurance | 10 |

### Test of Gross Motor Development (TGMD-2)

TGMD-2 [25] is used to assess the development of FMS in children aged 3-10 and has high validity and reliability[28–30]. It consists of a locomotor subtest (running, galloping, hopping, leaping, horizontal jumping, and sliding) and objects control subtest (striking a stationary ball, stationary dribbling, catching, kicking, overhand throwing and underhand rolling). Each subtest has 3-5 behavior components as a mature pattern of the skill. If a child performs correctly on the behavioral component, the rater marks 1, otherwise 0 is marked. The test was repeated twice by two examiners (by Pearson correlation coefficient test, r =0.63~0.81), and the average scores of two examiners was treated as the final result. Each of the original full score of the locomotor subtest and object control test was 48, where the highest is 96.

### FMS™

The FMS™ was conducted by the examiner with a certified FMS™ according to the standard protocol [19,31]. The test battery consisted of 12 test items including seven main tests: Deep Squat(DS), Hurdle Step (HS), in-line lunge (IL), Shoulder Mobility (SM), Active Straight Leg Raise(ASLR), Trunk Stability Push-Up(TSP), and Rotary Stability(RS), as well as three clearance tests: impingement clearing test, press-up clearing test and posterior rocking clearing test. Each test was listed with 3~5 specific action standards and the scores range from 0 to 3. If any pain occurred at any time during the testing, a score of 0 was given. In all tests except for the DS and TSP, both sides of the body were assessed separately. The lower score on both tests was recorded and counted into a final score. Seven tests score were compiled into a total of 21 points.

### Statistical Analysis

The original (raw) data obtained from the tests were input into Excel. Later, it was analyzed using SPSS version 25.0 for the analysis of the data. P value <0.05 was considered a significant difference for all tests. MF scores, TGMD-2 scores, and score FMS™ were tested for normal distribution of the data using Kolmogorov–Smirnov test. Descriptive statistics (mean ± SD) were used to express for all tests in each grade-group and sex-group. Kruskal-Wallis H test was used to determine differences in grade-group, and the Mann-Whitney U test was used to verify differences between gender groups. Pearson Correlation test was used to reveal correlation among the three groups of test results. Percentile cutoffs for both MF and FMS™ levels, good or high- (≥60th percentile), fair or moderate- (between the 36th and 59th percentiles), and poor or low- (≤35th percentile) were selected following Stodden [32]. Then the data were analyzed by multiple (3×3) chi-square tests of independence. For the first Chi-square analysis, FMS levels (T-A, T-B, T-C) were the independent variable, and MF levels (good, fair, or poor) were the dependent variable, and then FMS™ levels (high, moderate, or low) were taken as the dependent variable for the second chi-square analysis. Next, we changed the independent variable to FMS™ levels, and the MF levels, and FMS levels as the dependent variables for the twice (3×3) chi-square analysis, respectively. A *p-value* <0.05 was considered statistically significant.

## Results

### Basic situation analyses

#### Comparison of tests scores in different grades

Results of the MF test score (50.40±8.22), TGMD-2 score (68.36±8.46), and FMS™ (14.29±2.70) including responses of 117 children were carried out for the different groups respectively, and Kruskal-Wallis H test was used to compare the differences in mean. The results showed only the MF test and FMS™ had a significant difference in scores. Fig 2 presented the descriptive results of each grade group. In MF test, the significant difference result in 50 m sprint(*p*=0.035<0.05) and timed rope-skipping (*p* <0.001) (Fig 3).

**Fig 2.**
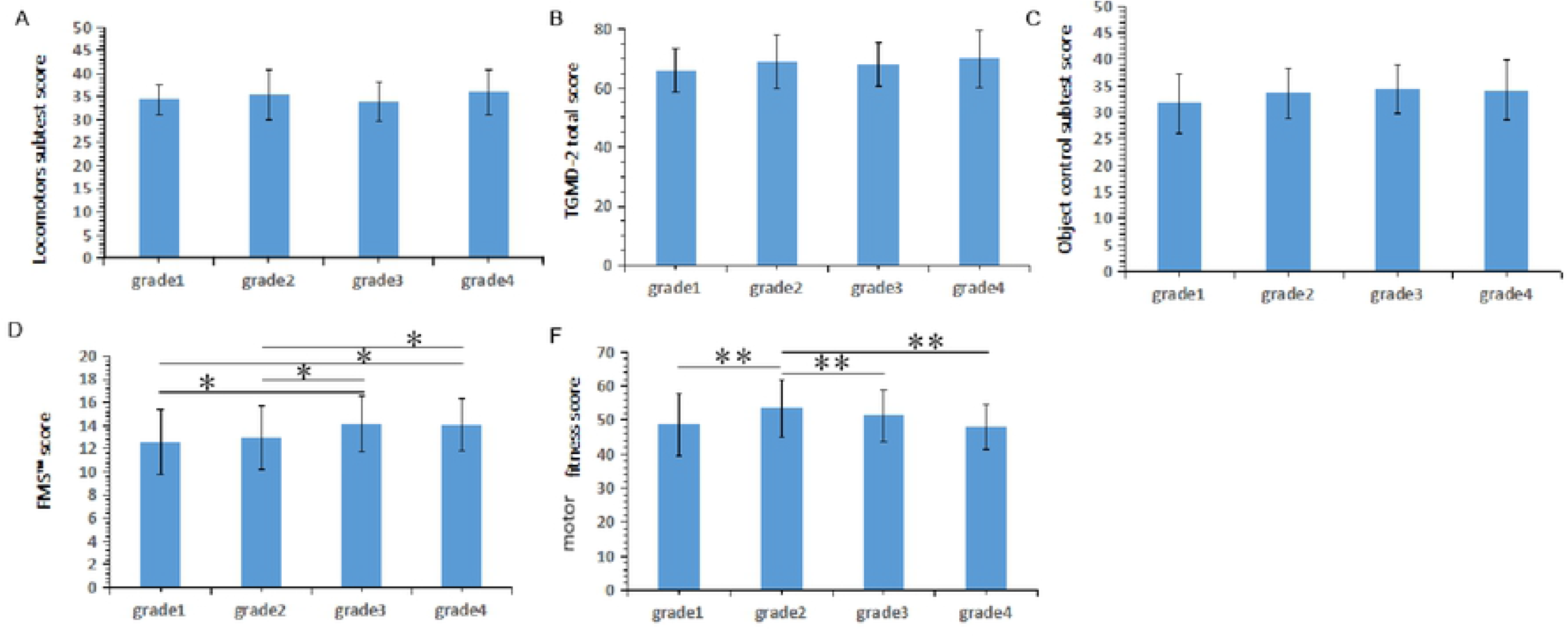
Descriptive statistics of TGMD-2, FMS™, and MF test from grade- groups (mean ± SD). *NOTE: TGMD-2 total score= Locomotors subtest score+ Object control subtest score* *Significant grade difference P < 0.05. ** Significant grade difference P < 0.01

**Fig 3.**
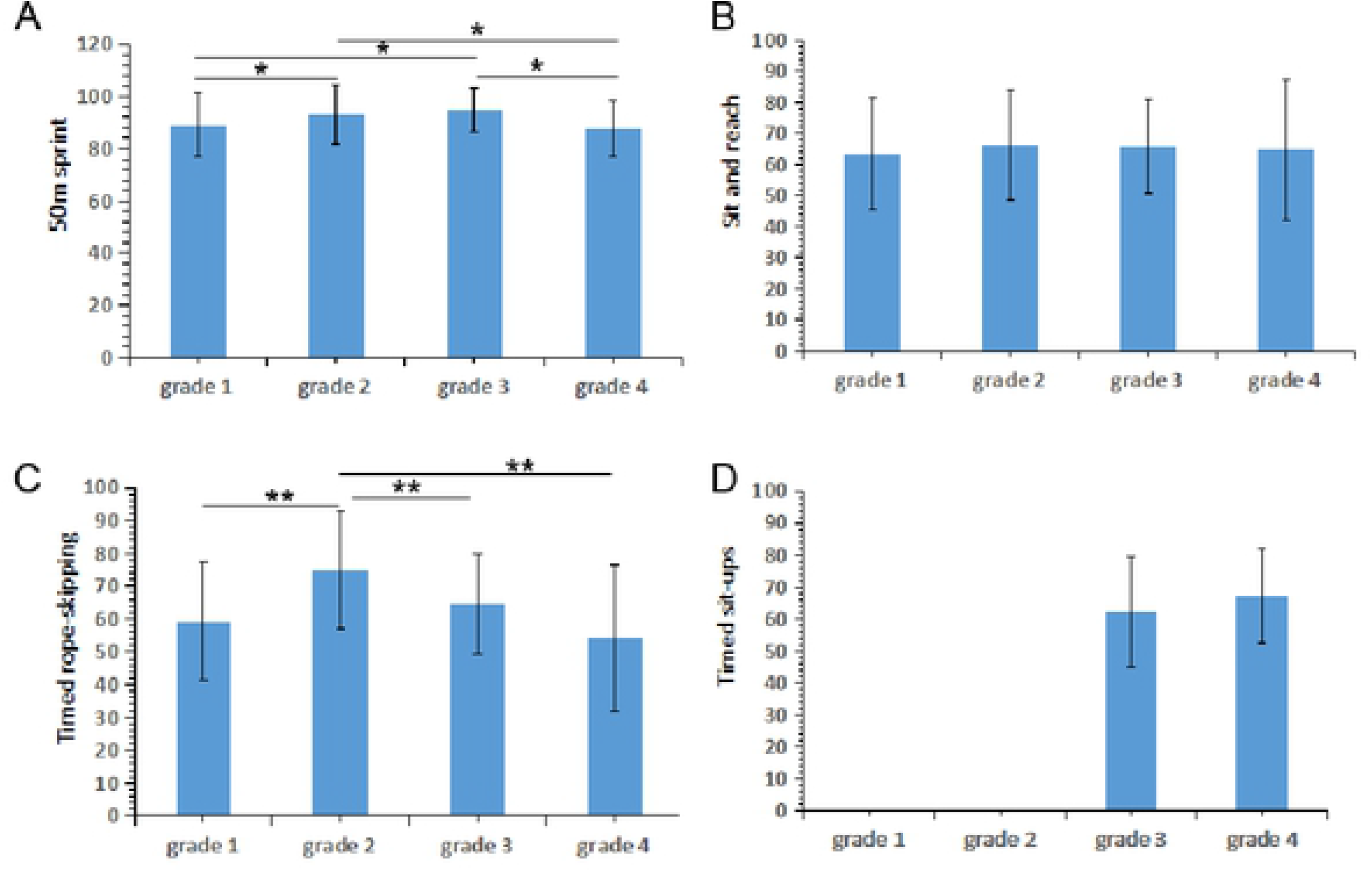
Descriptive statistics of subtest MF from grade- groups (mean ± SD). Note: MF results need to be weighed into the final MF test score. *Significant grade difference ***p*** < 0.05. ** Significant grade difference p<0.01

#### Comparison of scores in sex-group

Mann-Whitney U test was used to compare the difference in means in TGMD-2, FMS™, and MF test scores between sex groups. The boys scored significantly higher on the object control subtest score and the TGMD-2 total score; however, girls achieved a significantly higher score on FMS™ (p<0.001) (Table 3). Girls had higher scores than the boys concerning DS (p<0.01), HS (p=0.033), and ASLR (p<0.001). (Table 4).

**Table 3.** Descriptive statistics of TGMD-2, FMS™ and MF test between sex groups (mean ± SD).

|  | Locomotors | Object control | TGMD-2 | FMS<sup>TM</sup> | MF score |
| --- | --- | --- | --- | --- | --- |

|  | subtest score | subtest score | total score | score |  |
| --- | --- | --- | --- | --- | --- |
| **Boys** | 35.03±4.52 | 34.75±4.65 | 69.78±8.03 | 12.74±2.57 | 49.61±8.64 |
| **Girls** | 34.57±4.62 | 31.15±5.25 | 65.72±8.70 | 14.73±2.24 | 51.85±7.28 |
| **Z** | -0.815 | -3.541 | -2.457 | -4.112 | -0.1046 |
| **p** | 0.415 | 0.000* | 0.014* | 0.000** | 0.296 |

NOTE: TGMD-2 total score= Locomotors subtest score+ Object control subtest score
\*Significant gender difference $p<0.05$
\*\* Significant gender difference $p<0.01$

**Table 4.** Descriptive statistics of subtest FMS™ from sex- groups (mean ± SD).

|  | DS | HS | IL | SM | ASLR | TSP | RS |
| --- | --- | --- | --- | --- | --- | --- | --- |
| **boys** | 2.13±0.60 | 1.75±0.49 | 1.91±0.55 | 2.14±0.86 | 1.49±0.72 | 1.78±0.62 | 1.54±0.60 |
| **girls** | 2.54±0.55 | 1.95±0.44 | 2.05±0.44 | 2.37±0.77 | 2.22±0.73 | 1.85±0.52 | 1.76±0.58 |
| **Z** | -3.470 | -2.131 | -1.433 | -1.329 | -4.853 | -0.750 | -1.895 |
| **p** | 0.001* | 0.033* | 0.152 | 0.184 | 0.000* | 0.453 | 0.058 |

Note: DS=Deep Squat, HS=Hurdle Step, IL=in-line lunge, SM=Should Mobility,
ASLR=Active Straight Leg Raise, TSP=Trunk Stability Push-Up, and RS=Rotary
Stability
\* Significant gender difference $p<0.05$
\*\* Significant gender difference $p<0.01$

### The relevance of different levels of FMS groups with various levels of MF and FMS™

Using multiple 3×3 Chi-square tests analysis the relationship between FMS (TGMD-2 levels) and MF, no cell had expected count less than 5,χ^2^ (4,N=117) =14.605, *p* =0.006< 0.01, Cramer’s V = 0.25. These results indicate poor effect sizes (V = 0.25), thus suggesting that children with different FMS levels could have various MF levels, since there was a weak correlation between FMS and MF levels. Post hoc testing results are shown in Table 5. 38 participants had excellent FMS levels and were classified as the T-A group. It was noted that 60.5% were classified as “good” (≥ 60th percentile) MF levels, only 13% got the “poor” (≤ 35th percentile) MF levels, adjusted standard residuals >2, a significantly different probability than expected. When viewed from the low FMS skill participants, 53.8% in the T-C group demonstrated “poor” MF, and 16.7% showed a “good” MF level, unexpectedly both were significantly different. Therefore, these results may reflect the FMS incompetency as a “proficiency barrier” to Motor Fitness as discussed in Seefeldt’s developmental movement model.

**Table 5.** Chi-Square Cross-Tabulations for FMS Levels×MF Levels and FMS Levels×FMS™ Levels.

|  | <u>MF index levels</u> |  |  |  | <u>FMS™ index levels</u> |  |  |  |
| --- | --- | --- | --- | --- | --- | --- | --- | --- |
|  | Good | Fair | Poor | Total | High | Moderate | low | Total |
| T-A count | 23 | 9 | 6 | 38 | 23 | 9 | 6 | 38 |
| Expected count | 16.2 | 8.8 | 13.0 | 38 | 15.3 | 10.7 | 12 | 38 |
| % within | 60.5% | 23.3% | 15.8% | 100% | 60.5% | 23.7% | 15.8% | 100% |
| Adj.Std.Res. | (2.7) * | (0.1) | (-2.9)* |  | (3.1) * | (-0.8) | (-2.6)<br>* |  |
| T-B count | 18 | 9 | 13 | 40 | 11 | 12 | 17 | 40 |
| Expected count | 17.1 | 9.2 | 13.7 | 40 | 16.1 | 11.3 | 12.6 | 40 |
| % within | 45% | 22.5% | 32.5% | 100% | 27.5% | 30% | 42.5% | 100% |
| Adj.Std.Res. | (0.4) | (-0.1) | (-0.3) |  | (-2.0) * | (0.3) | (1.8) |  |
| T-C count | 9 | 9 | 21 | 39 | 13 | 12 | 14 | 39 |
| Expected count | 16.7 | 9 | 13.3 | 39 | 15.7 | 11.0 | 12.3 | 39 |
| % within | 23.1% | 23.1% | 53.8% | 100% | 33.3% | 30.8% | 35.9% | 100% |
| Adj.Std.Res. | (-3) * | (0) | (3.2) * |  | (-1.1) | (0.4) | (0.7) |  |
| Total count | 50 | 27 | 40 | 117 | 47 | 33 | 37 | 117 |
| Expected count | 50 | 27 | 40 | 117 | 47 | 33 | 37 | 117 |
| % within | 42.7% | 23.1% | 34.2% | 100% | 40.2% | 28.2% | 31.6% | 100% |

Note:\* Adjusted standard residuals (Adj. Std. Res.) statistically significant different probability than expected

The other 3×3 chi-square tests analysis the relationship between FMS (TGMD-2 levels) and FMS™, χ^2^ (4, N=117) =11.118, p =0.025 < 0.05, Cramer’s V = 0.218, also expressed FMS and have less relationship with FMS™. A similar phenomenon occurs in 60.5% children sorted as T-A group, which presented “high” FMS™ scores, and 15.8% showed the “low” scores. However, 33.3% of children in the T-C group, exhibited a “high” FMS™ score, and 35.9% got a “low” FMS™ score (Table 5).

### The relationship of levels of FMS™ groups with levels of MF and FMS

In the third Chi-square test (FMS™ levels × FMS levels) analysis, no cell had an expected count of less than 5. Additionally, χ2=11.118, p =0.025< 0.05, and Cramer’s V=0.218, indicated towards a similar trend. Through Post hoc testing shown in Table 6, we can see that 48.9% children classified as high FMS™ group, scored “well” on the FMS (T-A group); 27.7% showed “low” FMS ( T-C group). Likewise, only 16.2% of the “low” FMS™ group performed adequately in the TGMD test (T-A group), 45.9%, and 37.8% achieved B- and C- levels. In the fourth Chi-square analysis, the association of FMS™ levels and MF levels, no cell should have counted less than five, χ^2^=13.943 , *p* =0.007< 0.01, Cramer’s V=0.244. The chi-square analysis indicated 59.6% of “high” FMS™ children got a “good” score on the MF test, and 54.1% classified “low” FMS™ students, who presented “poor” MF performance.

**Table 6.** Chi-Square Cross-Tabulations for FMS™ Levels ×MF Levels and FMS™ levels ×FMS Levels.

|  | <u>MF index levels</u> |  |  |  | <u>FMS index levels</u> |  |  |  |
| --- | --- | --- | --- | --- | --- | --- | --- | --- |
|  | Good | Fair | Poor | Total | T-A | T-B | T-C | Total |
| High count | 28 | 9 | 10 | 47 | 23 | 11 | 13 | 47 |
| Expected count | 20.1 | 10.8 | 16.1 | 47 | 15.3 | 16.1 | 15.7 | 47 |
| % within | 59.6% | 19.1% | 21.3% | 100% | 48.9% | 23.4% | 27.7% | 100% |
| Adj.Std.Res. | (3.0) * | (-0.8) | (-2.4)* |  | (3.1) * | (-2.0) | (-1.1) |  |
| Moderate count | 13 | 10 | 10 | 33 | 9 | 12 | 12 | 33 |
| Expected count | 14.1 | 7.6 | 11.3 | 33 | 10.7 | 11.3 | 11.0 | 33 |
| % within | 39.4% | 30.3% | 30.3% | 100% | 27.3% | 36.4% | 36.4% | 100% |
| Adj.Std.Res. | (-0.5) | (1.2) | (-0.6) |  | (-0.8) | (0.3) | (0.4) |  |
| low count | 9 | 8 | 20 | 37 | 6 | 17 | 14 | 37 |
| Expected count | 15.8 | 8.5 | 12.6 | 37 | 12.0 | 12.6 | 12.3 | 37 |
| % within | 24.3% | 21.6% | 54.1% | 100% | 16.2% | 45.9% | 37.8% | 100% |
| Adj.Std.Res. | (-2.7) * | (-0.3) | (3.1) * |  | (-2.6)* | (1.8) | (0.7) |  |
| Total count | 50 | 27 | 40 | 117 | 38 | 40 | 39 | 117 |
| Expected count | 50 | 27 | 40 | 117 | 38 | 40 | 39 | 117 |
| % within | 42.7% | 23.1% | 34.2% | 100% | 32.5% | 34.2% | 33.3% | 100% |

Note: \* Adjusted standard residuals (Adj. Std. Res.) statistically significant different probability than expected

## Discussion

### Evaluation of motor performance of 7-10 years children

The first objective of this article was to compare MF, FMS, and FMS™ results of children aged 7-10 years. It is common for schools to have sports evaluations for children’s motor performance which ignores the evaluation of the process and quality of completed movements. In this study, we used the FMS™ and the TGMD-2 test, to compare the outcomes to those of the traditional MF test. The data showed that each grade of children’s MF score was not high. Furthermore, senior scores declined annually and significantly. The weak performance on timed sit-ups, sitting and reaching and timed rope-skipping reflects lack of flexibility, strength, and coordination.

From the TGMD-2 test, we found that locomotion subjects score (34.87±4.54) was significantly higher than the object control score (33.49±5.14, *p*=0.005<0.01). The total score of the fourth-grade children (70.13±9.68) was far from the full mature motor ability reported in the literature (full score: 96). Moreover, the score did not always increase with age, but fluctuated, indicating that the FMS of the school-age children from this sample source was at a moderate level. These results corroborate the results of another study [8] which assessed FMS proficiency of 6-9 years old children in Singapore and showed that most children’s locomotion skills were at “average” and “below average”, reaching “poor” and “below average” on control skills. Moreover, a survey [5] found low motor skill competency students were high in primary and high school students of Australia. It has also been shown [33] that only over 40% of children possessed proficiency in one set of skills (e.g., overhand throw in boys aged 10). Our findings suggest that motor development is related to age, but not entirely due to maturation.

Motor development is quite complex. We learn from Newell’s constraints model that skill acquirement depends on the interaction between constraints from an individual, environmental, and the task. [34] Most intervention studies and systematic reviews suggest that for children to learn motor skills they need t be provided with an accumulation of activity experience (i.e. participation in games and sports), formal instruction (i.e. physical education curriculum training[35–37] and PE teacher support. [38] Moreover, Tompsett [39] suggested that a combination of family practices seemed to improve FMS proficiency more effectively than school sports education alone.

A similar situation occurred in the case of FMS™ scores. The results showed that average scores (14.29±2.70) had a downward trend with age, half of 117 children scored less than or equal to 14, which supposed a cutoff value of injury prediction by restricted and asymmetrical fundamental movement patterns. Specifically, participants scored lowest in the TSP (1.80 ± 0.59) and RS (1.62 ± 0.6) items, indicating that school-age children had poor control of trunk strength and stability. It has been suggested [16] that correct movement patterns were originally formed through physical growth but which cannot maintain perfection because of weak or dysfunctional motor connection systems. Our results may be explained with the suggestion that due to increasing the load of studies with age, students have fewer opportunities to participate in exercises. Also, the adverse effects of family lifestyle may affect children (e.g. sedentary and high-calorie diets), which leads to the decrease of joint flexibility, stability, and coordination of movements, making fundamental movement patterns worse.

Furthermore, it is worth noting that motor performance had a gender difference in children. The results of the TGMD-2 test showed that boys’ object control skills exceeded that of girls, which was consistent with the rule of motor development. For example, boys’ overhand throw reached the mature stage when they were five years old, whereas girls in general only reached this stage after they were eight. The results of fundamental movement patterns showed that girls have significantly better scores than boys in DS, HS, and ASLR, which shows girls were stronger than boys in lower limb flexibility and stability of trunk. While teaching, we should pay attention to children’s gender differences and individual development. To strengthen core stability, boys’ flexibility and girls’ strength should be enhanced.

### Correlation between FMS scores, MF and FMS™

Our results show MF had a significant low positive correlation with FMS (p=0.006< 0.01, Cramer’s V = 0.25). These data also demonstrate that children with excellent FMS levels tend to score better in “good” MF (60.5%) and those who have bad FMS levels have less chances to score “good” on MF (16.7%). This evidence reflected the importance of the role of FMS in motor development. As has been suggested, [11] FMS provides the footstone for various physical activities in a pyramidal hierarchical model of motor development. If children cannot master more and wider FMS, they may have acquired a “proficiency barrier” to develop motor skills. [11] This has been investigated [32] in young adults (18-25y), and findings show low to moderate relevance among individual motor skill abilities and health-related fitness. There is possibly [40] a potential impact of low FMS on children with moderate-to-vigorous physical activity guidelines, providing the chance to fully develop children’s motor capacity. However, we should notice the small Cramer’s V value, which implies FMS do not completely predict MF.

We speculated that the progression of skill development is influenced by many factors, not just FMS, as it has been suggested [41] that physical and psychological factors may impede or promote skill development. Also, the traditional classifications of FMS may not be broad enough to encompass specific movement patterns, such as swimming strokes and push-ups.

Another unique aspect of our study was showing that MF also had positive correlation with FMS™ scores (*p* =0.007< 0.01, Cramer’s V=0.244), and FMS™ had a poor positive correlation with FMS (p =0.025 < 0.05, Cramer’s V = 0.218). The data showed high levels of FMS™ score can partially predict “good” MF (i.e.59.6%), and low levels of FMS™ scores of children with “good” MF was also less than the expected number (9 actual vs 15.8 expected). Further, 48.9% of children classified as high FMS™ group, had a “well” classification on FMS; those scoring low levels of FMS™, which was only a small sample, usually had the “well” classification on the FMS (i.e.16.2%), and others were more likely to have lower scores on the FMS test.

The concept of “FMS™” stems from the observation that babies learn basic movements in response to various stimuli. [42] We assumed FMS™ represented the basis of the Seefeldt’s pyramid. Our results showed FMS™, including reflexes and/or rudimentary movements, have a relationship with FMS and MF. Likewise, the “Performance Pyramid” suggests [43] the first level of function—movement—(i.e. FMS™) which represents flexibility and stability is appropriate to support other levels of function. On Cook’s “Performance Pyramid” model [43], the second layer depicts functional performance components, like endurance, strength, speed, and power, on top of is specific skill tests. It has also been suggested [44] that maintaining the correct movement pattern can optimize the quality of sports skill and reduce the consumption of energy, but there is still a low correlation Cramer’s V value between them. We can explain from the “Unskilled Performance Pyramid” [43] of existence that some people may have adequate functional movement patterns and efficiency of power generation, but also need supplementary training to master skills. It has been found [45] that four weeks of 30 minute fundamental movement training might affect specific isolated components of fitness, but not FMS™ performance.

Overall, our data revealed the lag of FMS development and degradation of functional movement pattern in primary school children. And although low, we also found a correlation between FMS™, FMS, and MF in the children’s sample. Results of post hoc testing demonstrated that children who had “good” functional movement patterns were more likely to be classified as “well” in the FMS, and those that are proficient in fundamental skills will tend towards efficient motor skills. Nevertheless, a relatively low correlation may indicate that these three links are relatively independent and influence each other, and they need to be integrated theoretically in a coherent fashion. In other words, at different periods of ontogeny or advantages of individual development, we may place much emphasis on developing one or a few aspects , meanwhile complementing and promoting each other. Lloyd and Oliver[46] provided a logical and physiological evidence-based “Youth Physical Development Model” (YPD), which showed that both FMS, physical ability(strength, agility, speed, etc.) and sport-specific skills are trainable at all times throughout childhood and adolescence (from age 2 to 21), but the emphasis placed on each component varies according to individual maturation[15].On the basis of such major findings and results, our future interventions need to be targeted at preschool level, which perhaps is the correct time-window for FMS competence improvement while allowing for healthy habits[5,47].

The strength of our study was using reliable tasks to examine the FMS hypothesis and functional movement patterns related to motor skills in primary school children. Our data can indicate some indirect evidence to the relationship among the three layers of Seefeldt’s pyramid model, and extend the associations between them, that is, three relatively independent systems linking with each other, generating mutual interaction and emerging throughout the whole process of motor development. When the development of motor skills hits a bottleneck, the relevant links should be re-examined and trained to finally achieve a breakthrough.

The study also has limitations. Because the TGMD-2 test norm is lacking in China, we chose the top, middle and bottom 10 TGMD-2 scores as a sample, which might have caused a gender gap in the sample. In our study boys outnumber girls by two to one, boys were selected because they scored higher in object control skills and high TGMD-2 total score.We also fixing percentile ranks to classify levels using “low, moderate, and high” and “good, fair, poor” terms, consistent with standard percentile rate data [32]. Due to the cross-sectional design of this study, we could not conclude a causality of the associations among the three. Rather, we explored a method and provided evidence to discuss how important these motor skills are for children. In the future, additional experimental and longitudinal studies are needed to use more accurate FMS assessments and permit studies to compare and improve the efficacy of FMS development [39]. Verifying the cause-and-effect relationship and mechanism between FMS and indicators related to children’s, even preschool children’s development, such as motor skills, physical fitness, and academic performance would also be necessary. Finally, it is noteworthy that we cannot ignore the influence social-ecological correlates (e.g., individual physical, psychological and social-cultural factors, nor the educational environment), which affects and restricts motor development [48–51].

## Acknowledgments

We would like to thank the primary school students for their cooperation and the teachers for their assistance. The experiment could not be carried out without them. The authors report no conflicts of interest.

